# Genomic analysis reveals limited hybridization among three giraffe species in Kenya

**DOI:** 10.1101/2023.09.12.557229

**Authors:** Raphael T. F. Coimbra, Sven Winter, Arthur Muneza, Stephanie Fennessy, Moses Otiende, Domnic Mijele, Symon Masiaine, Jenna Stacy-Dawes, Julian Fennessy, Axel Janke

## Abstract

**Background:** In the speciation continuum the strength of reproductive isolation varies, and species boundaries are blurred by gene flow. Interbreeding among giraffe (*Giraffa* spp.) in captivity is known and anecdotal reports of natural hybrids exist. In Kenya, Nubian (*G. camelopardalis camelopardalis*), reticulated (*G. reticulata*), and Masai giraffe sensu stricto (*G. tippelskirchi tippelskirchi*) are parapatric, and thus the country might be a melting pot for these taxa. We analyzed 128 genomes of wild giraffe, 113 newly sequenced, representing these three taxa.

**Results:** We found varying levels of Nubian ancestry in 13 reticulated giraffe sampled across the Laikipia Plateau most likely reflecting historical gene flow between these two lineages. Although comparatively weaker signs of ancestral gene flow and potential mitochondrial introgression from reticulated into Masai giraffe were also detected, estimated admixture levels between these two lineages are minimal. Importantly, contemporary gene flow between East African giraffe lineages was not statistically significant. Effective population sizes have declined since the Late Pleistocene, more severely for Nubian and reticulated giraffe.

**Conclusions:** Despite historically hybridizing, these three giraffe lineages have maintained their overall genomic integrity suggesting effective reproductive isolation, consistent with the previous classification of giraffe into four species.

## Background

Speciation is a continuous process that can be thought of as a spectrum of reproductive isolation [1]. Depending on the strength of the reproductive barriers, hybridization may lead to introgression and gene flow in areas of range overlap [2]. Introgressive hybridization can homogenize the genomic landscape of incipient species until they break down into hybrid swarms [3]. Alternatively, it may also enhance evolutionary potential by increasing the frequency of favorable genetic variants or introducing novel allele combinations [4]. These processes can create phylogenetic incongruence across the genome resulting in reticulate evolution and blurring species boundaries [5, 6]. Mounting evidence demonstrates that natural hybridization and gene flow between related species are common [7], as has been observed, for instance, in *Heliconius* butterflies [8], Darwin’s finches [9], and Grant’s gazelles [10].

Speciation and the number of species in giraffe (*Giraffa* spp.) has gathered considerable interest in recent years [11–17]. Giraffe have a wide and fragmented distribution throughout sub-Saharan Africa [18]. They are capable of long-distance movements up to 300 km [19] and can have home ranges as large as 1,950 km^2^ [20]. When housed together in captivity, some taxa can readily interbreed [21–23]. Yet, current taxonomic assessments based on nuclear and mitochondrial genetic data support either three [16] or four [17] highly divergent lineages with sub-structuring. Herein, we adopt the nomenclature used in Coimbra et al. [17], which includes four species and seven subspecies – the northern giraffe (*G. camelopardalis*), including West African (*G. c. peralta*), Kordofan (*G. c. antiquorum*), and Nubian giraffe (*G. c. camelopardalis* senior synonym of *G. c. rothschildi*); the reticulated giraffe (*G. reticulata*); the Masai giraffe sensu lato (*G. tippelskirchi*), including Luangwa (*G. t. thornicrofti*) and Masai giraffe sensu stricto (*G. t. tippelskirchi*); and, the southern giraffe (*G. giraffa*), including South African (*G. g. giraffa*) and Angolan giraffe (*G. g. angolensis*).

In East Africa, Nubian, reticulated, and Masai giraffe s. str. are largely parapatric [18] (Fig. 1a). In Kenya, their ranges adjoin in the seeming absence of natural barriers to dispersal in recent times [11, 24]. There have been anecdotal reports of individuals exhibiting intermediate phenotypes in the region [25–28], however, genetic admixture between giraffe species in the wild seems to be limited, implying that natural hybridization is rare [11, 14]. Reproductive asynchrony and seasonal variation in habitat use, both possibly related to regional differences in seasonality of rainfall and associated emergence and availability of browse, or pelage-based assortative mating may non-exclusively contribute to maintain genetic and phenotypic divergence (i.e., genetic structure in nuclear and mitochondrial DNA, and differences in pelage pattern) among these taxa [11, 24]. Nevertheless, hybridization between giraffe species in the wild has not been studied at a genomic scale.

**Fig. 1.**
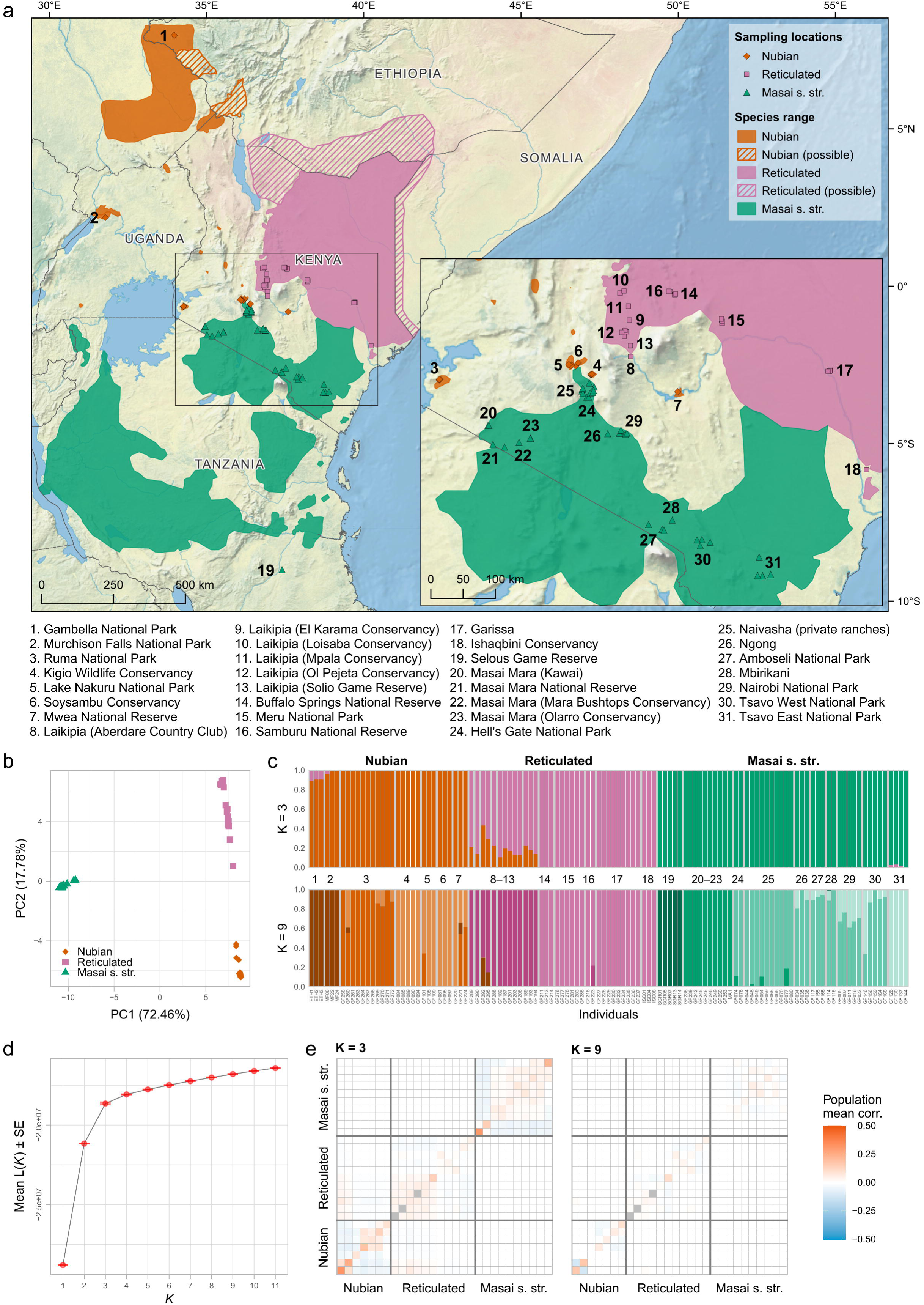
Population structure of Nubian, reticulated, and Masai giraffe s. str. (**a**) Geographical distribution of Nubian, reticulated and Masai giraffe s. str. (colored shadings) in East Africa and sampling locations (colored shapes and numbers). Hatched areas show estimated range of Nubian and reticulated giraffe populations. (**b**) PCA of 484,876 unlinked SNPs from 116 individuals representing Nubian, reticulated, and Masai giraffe s. str. PC1 separates Nubian and reticulated from Masai giraffe s. str., and PC2 separates Nubian from reticulated giraffe. The PCA space is further explored in Additional file 2: Fig. S2. (**c**) Ancestry proportions estimated from the same SNP dataset for *K* = 3 and *K* = 9. Colors indicate an individual’s cluster membership. The numbers in between plots represent sampling locations according to (a). Interspecies admixture is found mostly between Nubian and reticulated giraffe at *K* = 3 and is restricted to two individuals from Loisaba Conservancy at *K* = 9. Admixture analyses for *K* = 2–11 are shown in Additional file 2: Fig. S3. (**d**) Mean likelihood and standard error (SE) across 100 runs per *K*. Mean likelihoods start plateauing at *K* = 3. (**e**) Assessment of admixture model fit based on the correlation of residuals for *K* = 3 and *K* = 9. Plotted values are the mean correlation within and between individuals from each sampling locality. Model fit assessments for *K* = 1–11 showing the pairwise correlation of residuals between all individuals are available in Additional file 2: Fig. S4. The order of sampling localities is the same as in (c). Localities with only one sampled individual are shown in grey.

Contact zones provide unique opportunities to understand the nature of species boundaries and the processes involved in the onset and maintenance of speciation [29]. Moreover, as modern genomics enhances our ability to uncover species divergence in the presence of gene flow [30], our perception of hybridization and its consequences for the conservation of biodiversity deepens [31]. Here, we investigate the extent of hybridization and genetic admixture among the three giraffe taxa occurring in East Africa, focusing specifically on Kenya, and reconstruct changes in their population size in the recent past. We analyzed the complete nuclear and mitochondrial genomes of 128 wild giraffe sampled mostly across Kenya, including from suspected contact zones of Nubian and reticulated giraffe (i.e., the Laikipia Plateau; Fig. 1a, locations 8–13), and of reticulated and Masai giraffe s. str. (i.e., south of Garissa towards the Tsavo Region; Fig. 1a, locations 17, 30, and 31) [26]. This first genome-scale assessment of hybridization among East Africa’s giraffe lineages will aid to redefine their taxonomic status on the International Union for Conservation of Nature (IUCN) Red List and plan targeted conservation interventions for these threatened taxa [18].

## Results

### Genome resequencing

We analyzed genomes from 113 newly sequenced wild giraffe from across Kenya and 15 publicly available giraffe genomes from Ethiopia, Kenya, Tanzania, and Uganda (Fig. 1a and Additional file 1: Table S1), representing three separately evolving lineages: Nubian, reticulated, and Masai giraffe s. str. Read mapping against a chromosome-level Masai giraffe s. str. genome assembly [32] (GenBank: GCA_013496395) resulted in a mean mapping rate of 98.6% and a median filtered depth of 9× (1–26×) (Additional file 1: Table S1).

### Population structure and admixture

After filtering our dataset against relatedness (Additional file 2: Fig. S1), a principal component analysis (PCA) and an admixture analysis assuming three ancestry components (*K*) based on 484,876 unlinked single nucleotide polymorphisms (SNPs) correctly assigned all giraffe individuals to their respective species (Fig. 1). In the PCA (Fig. 1b), the first two principal components (PCs) explain most of the variance in the dataset, separating Nubian and reticulated from Masai giraffe s. str. (PC1: 72.46%), and Nubian from reticulated giraffe (PC2: 17.78%). On PC2, reticulated giraffe individuals from the Laikipia Plateau in Kenya (Fig. 1a, locations 8–13) are spread between Nubian and the remaining reticulated giraffe individuals. As we explore further PCs, they reveal population structure specific to each taxon (Additional file 2: Fig. S2).

In the admixture analysis (Fig. 1c and Additional file 2: Fig. S3), the plateauing of run likelihoods (Fig. 1d) and the residual fit of the admixture models (Fig. 1e and Additional file 2: Fig. S4) suggest that the number of *K* that better reflect the uppermost level of population structure in the data is *K* = 3. These three ancestry clusters correspond to the focal taxa of the study. As we increase *K*, the population structure within each taxon is revealed and we observe improvements in model fit up to *K* = 9 (Fig. 1c, Fig. 1e, and Additional file 2: Fig. S3 and Fig. S4). This indicates that the admixture model assuming *K* = 9 is the one that best explains the population structure in the data. In this model, most ancestry clusters correspond to groups of geographically close sampling localities. A notable exception is the cluster formed by Nubian giraffe from Gambella National Park (NP), in Ethiopia, and Murchison Falls NP, in Uganda – two locations which are geographically far apart. Individuals from these populations are grouped separately from each other in the PCA (Fig. 1b and Additional file 2: Fig. S2) and the residual fit of the admixture model for *K* = 9 shows a negative correlation between them (Fig. 1e, and Additional file 2: Fig. S4), suggesting that they have different population histories.

We detected signs of admixture from Nubian giraffe in 13 individuals (36.1%) of the reticulated giraffe between *K* = 3–5, with ancestry proportions at *K* = 3 ranging from 0.108–0.434. Like the observations in the PCA, these admixed individuals were all sampled in the Laikipia Plateau. At *K* ≥ 6, however, they are assigned to their own cluster with only two individuals from Loisaba Conservancy (GF292 and GF295) still showing signs of admixture from Nubian giraffe. In the Nubian giraffe, six individuals (19.4%) show admixture from reticulated giraffe between *K* = 3–4. Three of those individuals are from Gambella NP, with ancestry proportions at *K* = 3 ranging from 0.094–0.108, and three are from Murchison Falls NP, with ancestry proportions ranging from 0.002–0.038. However, at *K* = 5 these individuals form a separate cluster which seems to be the source of admixture of the 13 admixed reticulated giraffe individuals. Three individuals (6%) of Masai giraffe s. str. from Tsavo East NP also showed minimal admixture from reticulated giraffe at *K* = 3, with ancestry proportions from 0.023–0.035. However, at higher *K* values these proportions decrease approaching zero.

### Nuclear and mitochondrial phylogenies

We reconstructed maximum likelihood trees for two independent datasets: a set of 364,675 genome-wide SNPs from 125 giraffe, and a partitioned alignment of 13 mitochondrial protein-coding genes from 146 giraffe. For taxonomic completeness, both datasets included representatives of all four species and seven subspecies of giraffe with the okapi (*Okapia johnstoni*) as an outgroup. The tree topologies recovered (Fig. 2) are consistent with those reported in previous studies [14, 16, 17]. In the nuclear tree (Fig. 2a), individuals formed reciprocally monophyletic clades corresponding to their respective species with high support (UFboot ≥ 95 and SH-aLTR ≥ 80). Reciprocal monophyly of subspecies, however, was only supported for West African, Kordofan, and Nubian giraffe. Nubian and reticulated giraffe individuals that exhibited admixture signs in the ancestry clustering analysis are placed more externally within the clade of their respective taxa. In the mitochondrial tree (Fig. 2b), Luangwa and Masai giraffe s. str. cannot be distinguished, and the reticulated giraffe is paraphyletic. The grouping of Masai s. str. and South African giraffe is consistent with ancient mitochondrial introgression from Masai s. str. to South African giraffe, as reported in [15, 17], potentially representing a case of mitochondrial capture (i.e., complete replacement of the mitochondrial DNA of one species or population by another). Likewise, the observation of Masai giraffe s. str. individuals carrying reticulated giraffe mitochondrial haplotypes may indicate mitochondrial introgression from reticulated to Masai giraffe s. str., as suggested in [11], although incomplete lineage sorting (ILS) is also a plausible explanation.

**Fig. 2.**
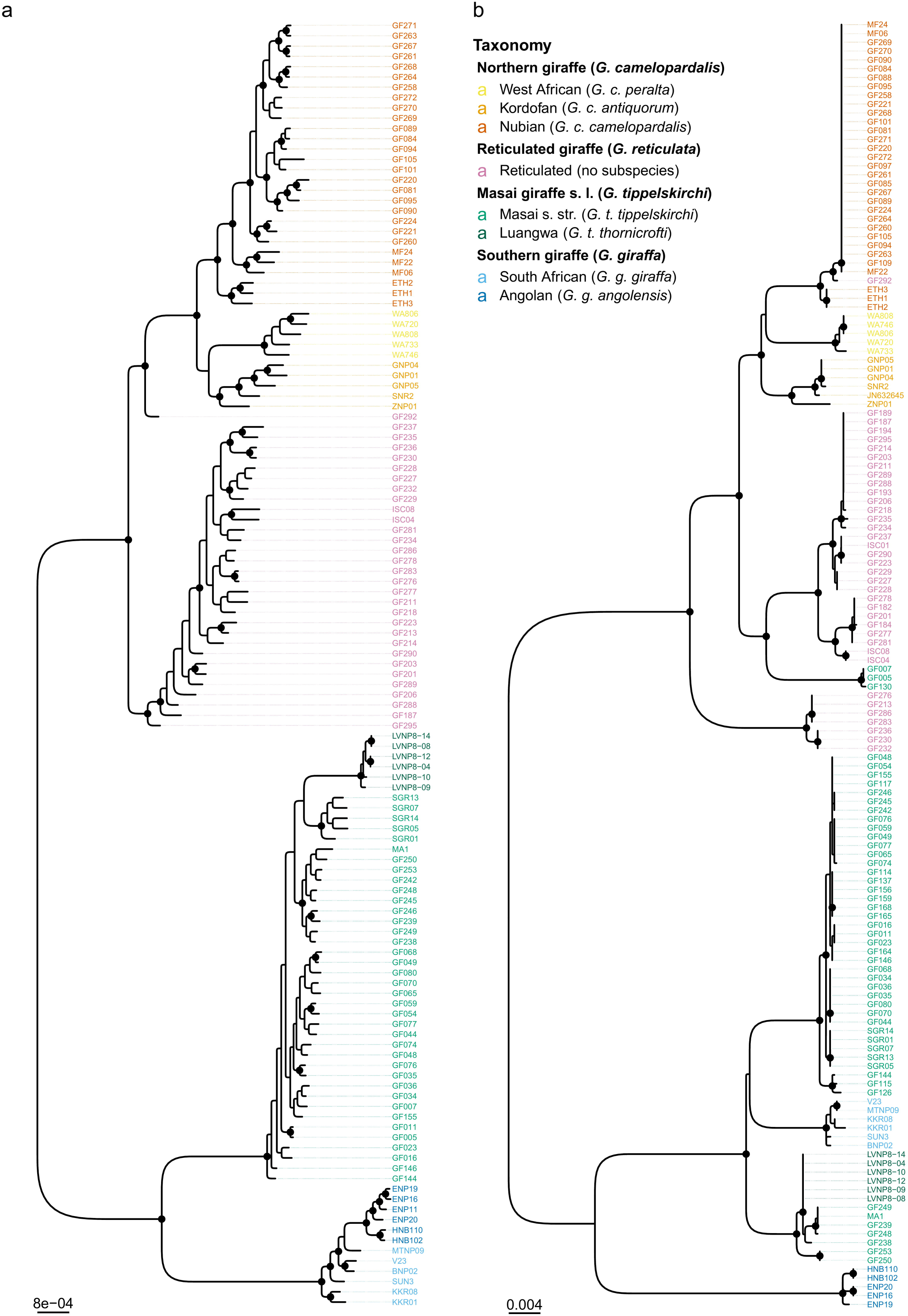
Nuclear and mitochondrial phylogenomic relationships among giraffe. Maximum likelihood phylogenies estimated from (**a**) 364,675 SNPs from 125 giraffe and (**b**) 13 mitochondrial protein-coding genes from 146 giraffe. The okapi was used as an outgroup (not shown). Colored tip labels indicate taxonomic assignment. Highly supported nodes (UFboot2 ≥ 95 and SH-aLTR ≥ 80) are marked with a black circle. In the nuclear tree, individuals formed clades corresponding to their respective species with high support. Mitochondrial introgression is observed from reticulated to Masai and from Masai to South African giraffe. Individual GF292 carries a Nubian giraffe mitochondrion and falls between the northern giraffe (i.e., West African, Kordofan, and Nubian) and the reticulated giraffe clades in the nuclear phylogeny; thus, likely representing a natural hybrid.

The individual GF292 carries a Nubian giraffe mitochondrion and falls between the northern giraffe (i.e., West African, Kordofan, and Nubian) and the reticulated giraffe clades in the nuclear phylogeny. That conforms with its high ancestry proportion from Nubian giraffe (0.434 at *K* = 3 and 0.299 at *K* = 9; Fig. 1c) and suggests that GF292 is either a recent reticulated × Nubian giraffe hybrid or more likely a backcross from a Nubian giraffe mother.

### Migration events and introgression

We estimated admixture graphs with migration events for Nubian, reticulated, and Masai giraffe s. str. populations (i.e., defined as the resulting clusters at *K* = 9) using the same dataset used for the SNP-based phylogenomic inference (Fig. 3a). Representatives of all four species and seven subspecies of giraffe were included for taxonomic completeness and the okapi was used as an outgroup. The topology of the estimated admixture graph is consistent with the SNP-based phylogeny. Further, an assessment of the optimal number of migration edges (*m*) allowed in the graph shows that one migration event (*m* = 1) is sufficient to explain over 99.8% of the variance in the data (Additional file 2: Fig. S5). However, including a second migration event (*m* = 2) improves the residual fit of the model (Additional file 2: Fig. S6). At *m* = 1, a migration event is modelled from Nubian giraffe to the reticulated giraffe from the Laikipia Plateau (locations 8– 13), while at *m* = 2, another migration is modelled from the branch leading to the reticulated giraffe clade to the base of the Masai giraffe s. l., albeit with a lower weight.

**Fig. 3.**
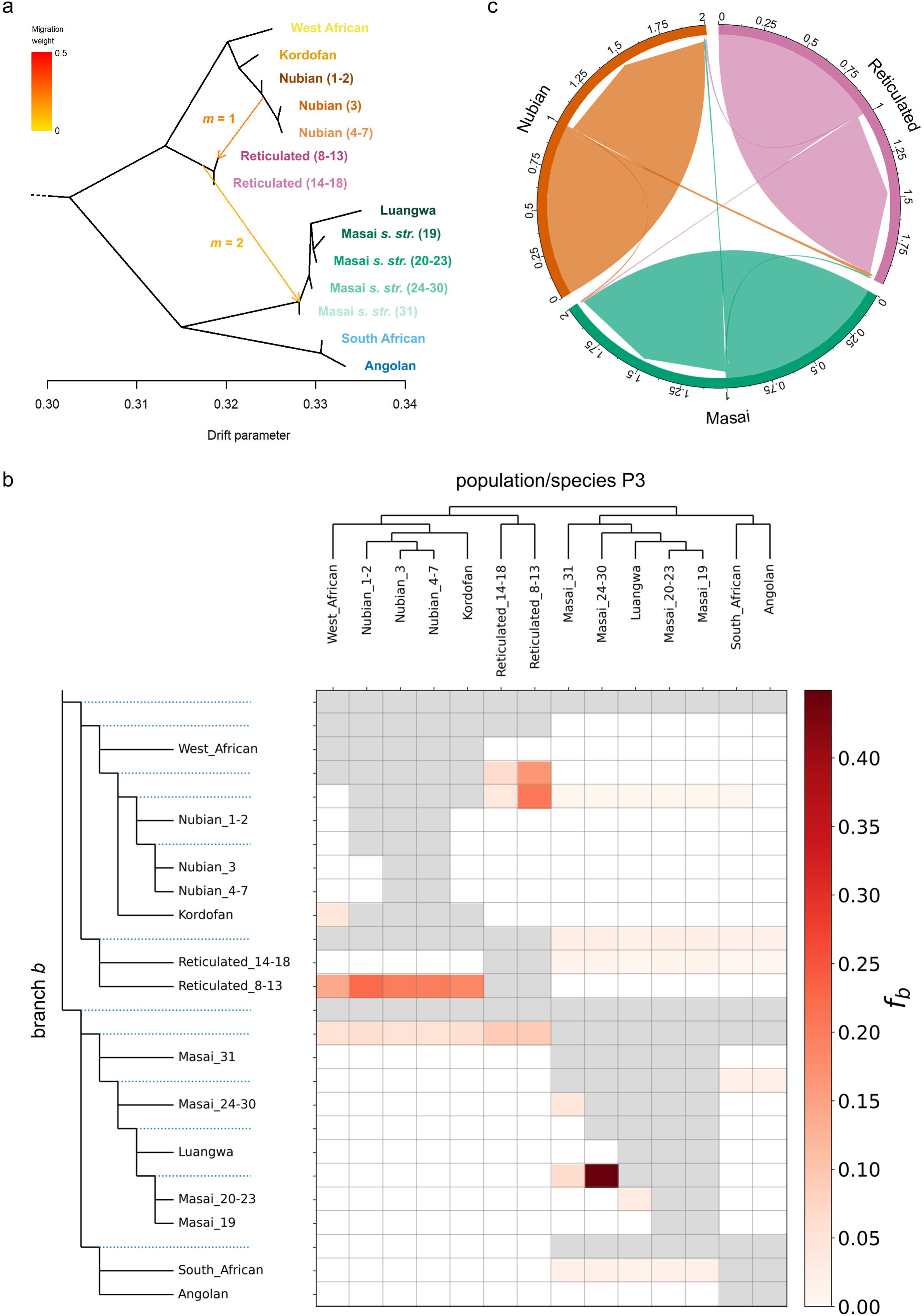
Signatures of gene flow among Nubian, reticulated, and Masai giraffe s. str. populations. (**a**) Admixture graph of giraffe populations with migration events. The okapi was used as an outgroup but was omitted from the image for a better resolution of the divergence between giraffe populations. Nubian, reticulated, and Masai s. str. giraffe populations were defined following the best fit admixture model (*K* = 9; Fig. 1c–e). Numbers within parentheses in Nubian, reticulated, and Masai s. str. giraffe population labels indicate sampling locations according to Fig. 1a. Migration arrows are colored according to their weight and marked following the number of migration events (*m* = 1 or *m* = 2) allowed in the model in which they were first inferred. The complete admixture graphs inferred for *m* = 1 and *m* = 2 and their corresponding residual fits are shown in Additional file 2: Fig. S6. (**b**) Heatmap showing the *f*-branch (*f*_b_) statistic estimated based on the topology recovered by OrientAGraph. *f*_b_ values are shown for tests where the *p*-value of the associated *D* statistic is < 0.01. Gray boxes indicate tips/branches which cannot be tested under a ((P1, P2) P3, Outgroup) topology. (**c**) Contemporary migration rates among Nubian, reticulated, and Masai giraffe s. str. Posterior mean migration rates were estimated based on 8,137 randomly sampled unlinked SNPs from 97 wild giraffe. Links with arrow tips indicate migration direction. Link widths are proportional to the fraction of individuals in the recipient population with ancestry in the source population (per generation). Scale ticks represent the cumulative fraction of migrants (per generation). Posterior estimates and 95% credible sets for migration rates are provided in Additional file 3: Table S2.

The inferred admixture graph topology was used as a guide tree to calculate the *f*-branch (*f*_b_) statistic [33] based on genotype probabilities from the same SNP dataset. That was done for all possible giraffe population trios using the okapi as an outgroup. The *f*_b_ assigns evidence for introgression (i.e., *f*_4_-ratio [34] scores) to specific branches on a population/species tree, including internal branches, thus conveying information about the timing of introgression. In our analysis, the *f*_b_ identifies a total of 48 signals of excess allele sharing between the population/species P3 (Fig. 3b, x-axis) and the branch *b* (Fig. 3b, y-axis); however, *f*_b_ ≥ 0.05 in only 17 of them. In particular, the *f*_b_ signals suggest gene flow events between reticulated giraffe from Laikipia Plateau (locations 8–13) and the branches leading to Kordofan + Nubian giraffe (*f*_b_ = 0.16) and to Nubian giraffe populations (*f*_b_ = 0.21). Weaker *f*_b_ signals also indicate gene flow between reticulated giraffe populations (locations 8–13 and 14–18) and the branch leading to Masai s. str. + Luangwa giraffe (*f*_b_ = 0.09 in both cases). However, the strongest identified *f*_b_ signal (*f*_b_ = 0.45) corresponds to gene flow between Masai giraffe s. str. populations from locations 24–30 (i.e., Amboseli NP, Hell’s Gate NP, Mbirikani, Nairobi NP, Naivasha, Ngong, and Tsavo West NP) and the branch leading to Masai giraffe s. str. populations from the Selous (location 19) and Masai Mara Game Reserves (locations 20–23). In all those cases, gene flow from P3 (x-axis) into branch *b* (y-axis) also generated horizontal lines of correlated *f*_b_ signals between branch *b* and lineages related to P3 due to their shared ancestry with P3 [35].

### Contemporary gene flow

We estimated both directionality and rates of contemporary migration (last two generations) between Nubian, reticulated, and Masai giraffe s. str. based on a subset of 8,137 unlinked SNPs from 97 individuals with median read depth ≥ 8. The highest mean posterior migration rate is observed from Nubian to reticulated giraffe, where 2% of the individuals in the reticulated giraffe are estimated to be migrants derived from the Nubian giraffe (per generation) (Fig. 3c and Additional file 3: Table S2). Migration rates inferred between other species in any direction are ≤ 1.1%. However, all 95% credible intervals for migration rates include zero, and therefore absence of recent gene flow cannot be statistically rejected (Additional file 3: Table S2).

### Demographic reconstruction

Reconstruction of population size changes over the recent past based on the site frequency spectrum (SFS) reveals a general decrease in effective population sizes (*N_e_*) for the three analyzed giraffe taxa (Fig. 4). We observe similar but unsynchronized demographic trends with an accentuated population bottleneck (Nubian: ∼6.5–18 ka ago; reticulated: ∼28–54 ka ago, Masai s. str.: ∼10.5–25 ka ago) intercalating periods of relative stability at higher *N_e_* (Nubian: ∼0.7–5 ka and ∼18–76 ka ago; reticulated: ∼3–20 ka and ∼54–140 ka ago, Masai s. str.: ∼2–7 ka and ∼25–60 ka ago). The Nubian and the reticulated giraffe experienced a sharp decline between ∼0.48–0.7 ka and ∼2.5–3 ka ago, respectively, towards an approximately constant *N_e_*, while the Masai giraffe s. str. shows a gradual decline between ∼0.7–2 ka before reaching relative stability. Population bottlenecks older than 50 ka are observed for all three giraffe taxa; however, these cannot be interpreted reliably due to limitations of SFS-based demographic methods for ancestral time spans [36]. Overall, median *N_e_* dropped from their highest ancestral estimates of ∼62,400 to currently ∼2,700 for the Nubian giraffe, from ∼130,200 to a present ∼5,500 for the reticulated giraffe, and from ∼29,500 to ∼1,700 for the Masai giraffe s. str.

**Fig. 4.**
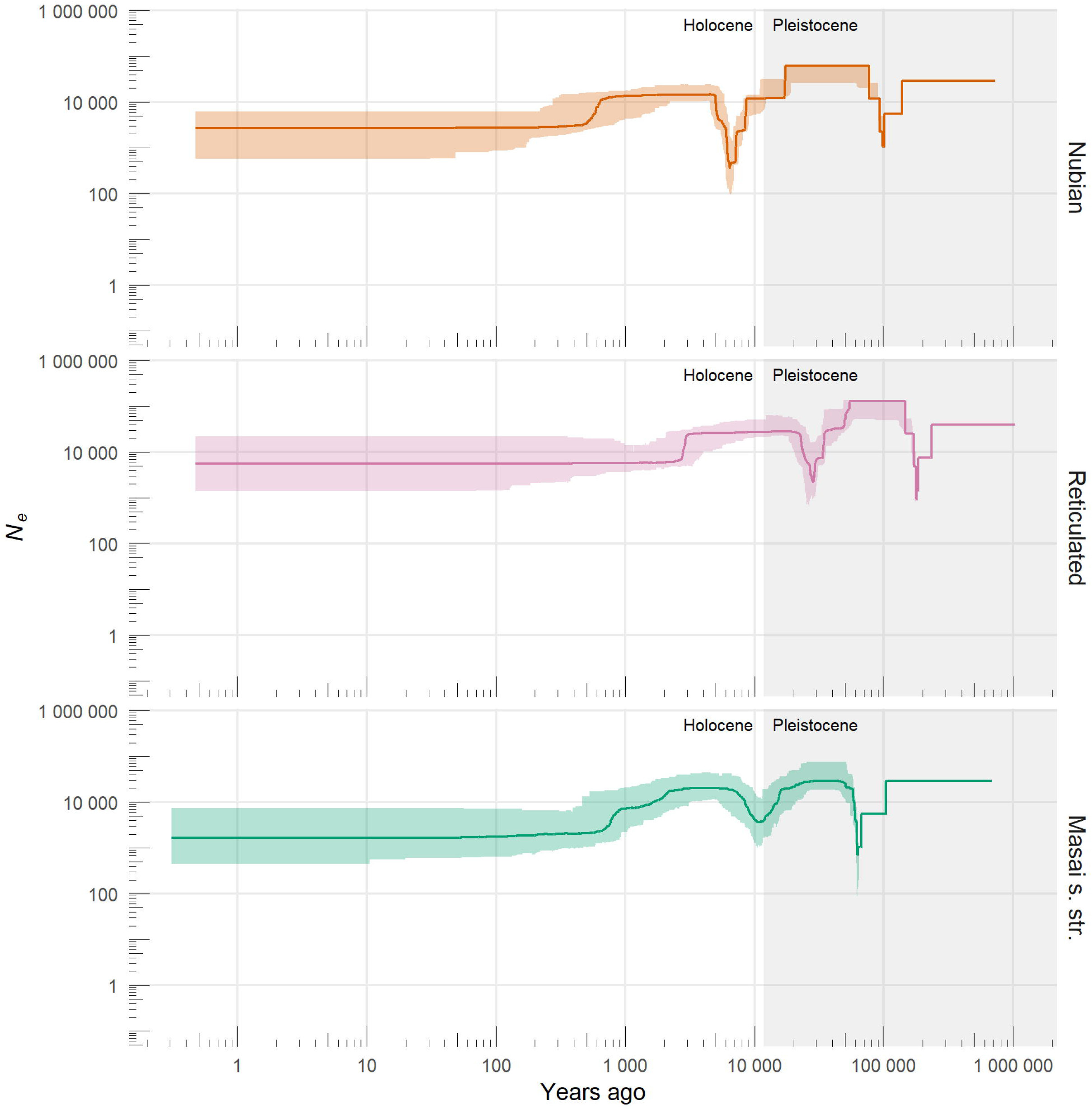
Demographic history of Nubian, reticulated, and Masai giraffe s. str. Population size changes over the recent past were reconstructed from the site frequency spectrum (SFS) using the stairway plot model after masking singletons. Axes were scaled by a mutation rate of 2.12 × 10^-8^ substitutions per site per generation and a generation time of 10 years. Colors represent focal giraffe taxa. Solid lines indicate median *N_e_* estimates and shaded areas correspond to 95% confidence intervals.

## Discussion

The number of giraffe species has been a subject of debate, particularly the question whether northern and reticulated giraffe should be considered separate species [14–17]. As the ‘species’ is predominantly used as the fundamental unit of conservation and as a metric of biodiversity [37], understanding species distinction is key for accurate conservation status assessments that can effectively guide conservation efforts [18]. In addition, failing to identify or neglecting admixed populations and hybrids can be detrimental to conservation policy making, and thus to species conservation [38]. Our findings corroborate significant genetic divergence between northern, reticulated, and Masai giraffe s. l., as shown by the distinct PCA and admixture clusters, as well as their reciprocal monophyly in the SNP phylogeny. In the mitochondrial tree, the nesting of northern giraffe within a paraphyletic reticulated giraffe may be explained by ILS resulting from peripatric or “budding” speciation [39] in peripheral populations of reticulated giraffe. However, under peripatric speciation, parallel patterns of paraphyly are expected across nuclear and mitochondrial loci [39], and this was not observed in our dataset. Furthermore, while three Masai giraffe s. str. individuals were identified carrying reticulated giraffe mitochondria, we cannot confidently distinguish between ancient mitochondrial introgression and/or ILS as the likely cause.

As demonstrated, while introgressive hybridization between Masai giraffe s. str. and other giraffe taxa in Kenya is minimal, admixture between Nubian – the easternmost subspecies of the northern giraffe – and reticulated giraffe seems to be asymmetrical towards the latter and restricted to a contact zone in the Laikipia Plateau. Although hybridization among giraffe in that region has been previously conjectured, with the observation of individuals exhibiting intermediate phenotypes (i.e., pelage pattern) [26, 28], we provide the first genomic evidence of its occurrence. This finding reinforces the unreliability of morphological characters such as pelage pattern for the identification of giraffe (sub)species, especially at a local scale [27]. Moreover, while the estimated migration events in the admixture graph and the *f*_b_ statistic support gene flow between Nubian and reticulated giraffe across the Laikipia Plateau, they also suggest that such an event is most likely ancestral. Consistent with that, estimates of contemporary migration rates were low and not statistically significant suggesting that current gene flow is limited. In fact, only two reticulated giraffe individuals consistently show signs of admixture from Nubian giraffe upon deeper investigation of population structure at *K* ≥ 6. Habitat loss and fragmentation, linked with human population growth, has drastically reduced the natural range of the Nubian giraffe in Kenya, such that most of its present-day populations are a result of extralimital translocations from a source population near Eldoret during the 1970s [18, 40]. All these introductions were into government or private fenced wildlife areas, which would have further restricted gene flow between Nubian and reticulated giraffe over the last few generations. More importantly, however, the overall genomic integrity of the parent taxa despite the existence of a contact zone suggests reproductive isolation and can be interpreted as support for species status [41].

Previous studies hypothesized that the divergence between giraffe lineages in East Africa could be linked to climatic oscillations associated with Earth’s precession cycles during the Pleistocene [11, 24]. Divergence could have been triggered either by spatially and temporally contrasting rainfall regimes that persisted until the Early Holocene, or by repeated expansion and contraction of the savannah habitat resulting in periodic isolation in refugia [24]. Nevertheless, the long-term maintenance of genetic distinctiveness between them appears to correlate with regionally distinct rainfall patterns in East Africa and the associated differences in timing of emergence and availability of browse [24]. Reproductive asynchrony resulting from potential local adaptation of the reproductive cycle to the differential timing of green-up may explain such correlation [11, 24]. That would imply that hybrids may display reduced fitness if they were born in an unfavorable season [11, 24], and result in negative selection against them, which would likely restrain introgression to contact zones [41]. Introgressive hybridization from Nubian into reticulated giraffe in Laikipia may have occurred under these conditions. Nubian giraffe populations have been shown to exhibit temporal and seasonal migration patterns [42]. In our study, this is particularly important considering the proximity and lack of a geographic barrier between the Lake Baringo Basin, a historical Nubian giraffe stronghold, and the Laikipia Plateau, which currently holds ∼28% of the extant reticulated giraffe population. Conversely, the absence of substantial admixture involving Masai giraffe s. str. could reflect stronger selection against its hybrids. Seasonal variation in habitat use (i.e., resource tracking) and pelage-based assortative mating could also affect the maintenance of genetic divergence and broad-scale phenotypic differences between the taxa [11, 24].

Our reconstruction of demographic changes for the three focal taxa in the recent past expands previous inferences made for the distant past [17], and provide a more complete picture of their population history. In general, estimates of *N_e_* for the three giraffe lineages were higher during the Late Pleistocene than they are today. Population reductions observed since the Late Pleistocene–Holocene transition could be linked to a period of wetter conditions and associated contraction of savannahs that lasted from ∼5.5–14.8 ka ago [43], and later to the spread of pastoralism in East Africa from ∼4.7 ka ago onwards [44]. *N_e_* values estimated at the present are reasonable, and expectedly lower [45] than the current estimated census population sizes (*N_c_*) [18]. The reticulated giraffe has the highest present-day *N_e_* (∼5,500) among the three taxa, while the Masai giraffe s. str. has the lowest (∼1,700). Furthermore, the Masai giraffe s. str. shows the largest difference between present-day *N_e_*(∼1,700) and *N_c_* (∼44,700 [18]), followed by the reticulated giraffe (*N_e_* = ∼5,500 and *N_c_* = ∼16,000 [18]), and the Nubian giraffe (*N_e_* = ∼2,700 and *N_c_* = ∼3,000 [18]). These observations are in line with previous findings of genomic diversity that is higher in reticulated, moderate in Nubian, and lower in Masai giraffe s. str. [17].

### Conclusions

Our findings reinforce the classification of giraffe into the four species (i.e., northern, reticulated, Masai s. l., and southern giraffe) proposed in [13, 14, 17], by clearly separating northern from reticulated giraffe, with limited recent introgression reflecting effective reproductive isolation. These results have valuable and direct conservation implications for the management of giraffe in Kenya and more broadly throughout their range in Africa. As the three species present in Kenya are genetically distinct, it is important that future conservation interventions, such as translocations, take these findings into account to avoid mixing distinct (sub)species, hence maintaining their unique biodiversity [46]. The outcome of this study is critical to appropriate re-classification of giraffe on the IUCN Red List and in turn informing targeted conservation actions for each taxon, particularly for African range states and international convention reviews such as the Convention on International Trade in Endangered Species of Wild Fauna and Flora (CITES) [47, 48]. Moreover, the comprehensive genomic dataset made available here constitutes an important resource for future studies of local adaptation in the giraffe lineages in East Africa. By identifying loci under selection and deeply characterizing the genetic composition of admixed/hybrid individuals, it might be possible to shed light on the potential effects of such loci on the fitness of giraffe hybrids in nature.

## Methods

### Sampling

Skin biopsy samples from 113 wild giraffe (Nubian, *n* = 32; reticulated, *n* = 37; Masai s. str., *n* = 44) from different parts of Kenya were collected as a collaboration between the Giraffe Conservation Foundation (GCF) and Kenya Wildlife Service (KWS), together with local partners, via remote biopsy darting and carcasses, and preserved in absolute ethanol. Sampling was conducted with the appropriate access and research permits from the Kenyan authorities. Sampling locations and sample details are presented in Fig. 1a and Additional file 1: Table S1. Short reads from wild individuals of West African (*n* = 5), Kordofan (*n* = 5), Nubian (*n* = 6), reticulated (*n* = 3), Masai s. str. (*n* = 6), Luangwa (*n* = 6), South African (*n* = 6), and Angolan giraffe (*n* = 6) analyzed in Coimbra et al. [17] and Agaba et al. [49] were added to the new dataset for a comprehensive representation of all four species and seven subspecies of giraffe (Additional file 1: Table S1). The okapi from Agaba et al. [49] was included in analyses that required an outgroup.

### Whole-genome sequencing

DNA was extracted using either a NucleoSpin Tissue Kit (Macherey-Nagel) or the phenol-chloroform protocol [50]. Sequencing libraries were prepared with the NEBNext Ultra II DNA Library Prep Kit (New England Biolabs, Inc.) at Novogene and sequenced on an Illumina NovaSeq 6000 (2 × 150 bp, 350 bp insert size).

### Read processing

Quality control of short reads was done in fastp v0.20.0 [51] with base correction and low complexity filter enabled. Adapters and polyG stretches in read tails were automatically detected and removed. Trimming was performed in a 4-bp sliding window (option --cut_right for reads from Agaba et al. [49] and --cut_tail for the remaining) with a required mean base quality ≥ 15. Reads shorter than 36 bp, composed of > 40% of bases with quality < 15, or containing > 5 Ns were discarded.

### Read mapping

Reads were mapped against a chromosome-level Masai giraffe s. str. genome assembly [32] (GenBank: GCA_013496395) with BWA-MEM v0.7.17-r1188 [52]. The resulting SAM files were coordinate-sorted, converted to BAM, and merged to sample-level with samtools v1.10 [53]. Duplicate reads in the BAM files were marked with MarkDuplicates from Picard v2.21.7 (http://broadinstitute.github.io/picard/) and regions around indels were realigned with GATK v3.8.1 [54]. Reads mapped to repetitive regions identified with RepeatMasker v4.0.7-open [55] and to sex chromosomes, unmapped reads, and reads flagged with bits ≥ 256 were removed from the BAM files with samtools. Only reads mapped in a proper pair were retained.

### SNP calling and linkage pruning

SNPs were called per species in Nubian, reticulated, and Masai giraffe s. str. individuals with median read depth ≥ 2 (Additional file 1: Table S1). Genotype likelihoods were estimated in ANGSD v0.933 [56] with options -GL 1 -baq 2. Minimum mapping and base quality scores were set to 30 and depth thresholds were set to *d* ± (5 × *MAD*), where *d* is the median and *MAD* is the median absolute deviation of the global site depth distribution. Only biallelic SNPs called with a *p*-value < 1 × 10^-6^, present in at least 90% of the individuals, and with a minor allele frequency (MAF) of at least 5% were considered. SNPs were tested for strand bias, heterozygous bias, and deviation from Hardy-Weinberg equilibrium (HWE) and discarded if any of the resulting *p*-values was below 1 × 10^-6^.

Each species’ SNP set was then independently pruned for linkage disequilibrium (LD) with ngsLD v1.1.1 [57]. We estimated *r^2^* values for SNP pairs up to 500 kbp apart as a proxy for LD and plotted linkage decay curves per species from a random sample of 0.05% of the estimated *r^2^* values (Additional file 2: Fig. S7). Appropriate thresholds for linkage pruning were selected based on each species’ linkage decay curve. SNPs were pruned assuming a maximum pairwise distance of 100 kbp for all species and a minimum *r^2^* of 0.10 for reticulated and Masai giraffe and 0.15 for the Nubian giraffe. The resulting pruned SNPs from each species were concatenated and used as input in a second SNP calling round to generate a combined dataset with individuals from the three species. SNPs were jointly called in ANGSD with the -sites option and no MAF, HWE, or SNP *p*-value filters were used. All remaining settings were as described above.

### Relatedness

The combined SNP dataset generated above was used to estimate pairwise relatedness in NGSremix v1.0.0 [58], which accounts for individuals with admixed ancestry. NGSremix additionally requires admixture proportions and ancestral allele frequencies as inputs, which were obtained from the run with the highest log-likelihood out of 100 runs of NGSadmix v32 [59] assuming three ancestry components (*K*). A custom R script was used to identify and select closely related individuals that, when removed from the dataset, would maximize the reduction of the overall relatedness in the data while minimizing sample loss (Additional file 1: Table S1 and Additional file 2: Fig. S1). These individuals were removed from all further analyses. A final round of joint SNP calling with ANGSD was performed as described above to obtain a combined SNP dataset for Nubian, reticulated, and Masai giraffe s. str. that was LD pruned and filtered against relatedness.

### Population structure and admixture

Genotype likelihoods of unlinked SNPs estimated in ANGSD were used to calculate a covariance matrix in PCAngsd v1.03 [60]. A PCA was then performed using the prcomp() function in R v4.2.2 [61]. Ancestry clusters and individual ancestry proportions were inferred in NGSadmix assuming a *K* value ranging from 1 to 11. The analysis was repeated 100 times per *K* with different random seeds and the replicate with the highest log-likelihood for each *K* ≥ 2 was shown as an admixture plot. The fit of the resulting admixture models was assessed based on the pairwise correlation of residuals between individuals estimated with evalAdmix v0.962 [62].

### SNP-based phylogenomic inference

Genotypes were jointly called in individuals from all giraffe species and subspecies with median read depth ≥ 8 with the okapi as an outgroup (Additional file 1: Table S1). Genotype calling was performed using bcftools v1.17 [53] mpileup + call pipeline, with option --full-BAQ and minimum mapping and base quality set to 30. Samples were grouped per (sub)species (--group-samples) and HWE assumption was applied within but not across groups. The commands filter and view were used to convert genotypes with GQ < 20 to missing data and filter for biallelic SNPs with a fraction of missing genotypes ≤ 0.1, QUAL ≥ 30, MQ ≥ 30, and within depth thresholds calculated as described for ANGSD. To reduce the impact of LD, we randomly sampled ∼1% (462,697) of all SNPs using vcfrandomsample from vcflib v1.0.3 [63]. The called genotypes from the subsampled VCF were used create a matrix for phylogenetic analysis in PHYLIP format with vcf2phylip v2.8 [64]. After removing constant, partially constant, and ambiguously constant sites from the SNP matrix, a maximum likelihood phylogeny was constructed in IQ-TREE v2.2.2.3 [65] based on 364,675 SNPs. Ultrafast model selection [66] between nucleotide substitution models with ascertainment bias correction [67] was enabled. Branch support was assessed by 1,000 replicates of the ultrafast bootstrap approximation (UFBoot) [68], with hill-climbing nearest neighbor interchange (NNI) optimization, and the Shimodaira–Hasegawa-like approximate likelihood ratio test (SH-aLRT) [69]. The tree was plotted with ggtree v3.6.2 [70].

### Assembly and phylogeny of mitochondrial genomes

Mitogenomes were assembled *de novo* from the unprocessed reads using GetOrganelle v1.7.4 [71] with options –-fast -k 21,45,65,85,105 -F animal_mt. In some instances, fine-tuning parameters -w and –max-n-words was necessary to obtain a complete circular genome. Mitogenome assembly sequences were visually inspected and curated (i.e., corrected directionality, circularized, adjusted starting nucleotide) on Geneious Prime v2020.1.2 (https://www.geneious.com/).

Sequences of all 13 mitochondrial protein-coding genes were extracted from the assemblies and aligned to sequences of wild giraffe analyzed in Coimbra et al. [17] and Hassanin et al. [72] (GenBank: JN632645) with the L-INS-i algorithm in MAFFT v7.475 [73]. The okapi (GenBank: JN632674) [72] was also included as an outgroup for phylogenetic inference. A maximum likelihood phylogeny was constructed in IQ-TREE through a partitioned analysis [74] of the protein-coding gene alignments. Ultrafast model selection between codon models was enabled assuming the vertebrate mitochondrial genetic code. Branch support was assessed with 1,000 replicates of the UFBoot and the SH-aLRT. The tree was plotted with ggtree.

### Inference of migration events

The topology and migration events between the Nubian, reticulated, and Masai giraffe s. str. populations (defined as the clusters resulting from the best fitting admixture model) were inferred as admixture graphs with TreeMix v1.13-r231 [75] and OrientAGraph v1.0 [76]. The subsampled VCF generated for the SNP-based phylogenomic inference was processed with PLINK v1.9 [77] and plink2treemix.py (https://bit.ly/3LCcNW4) to obtain allele counts per population as input. TreeMix was ran using blocks of 100 SNPs and assuming the number of migration edges (*m*) ranging from 0 to 5 for 50 independent optimization runs per *m*. The okapi was used to root the graph. A range of *m* values to be further explored was determined by looking at the Δ*m* and the percentage of explained variance estimated with OptM v0.1.6 [78]. We then ran OrientAGraph with options -mlno -allmigs for 10 bootstrap replicates assuming the selected *m* values. OrientAGraph improves TreeMix’s heuristics with an exhaustive search for a maximum likelihood network orientation (MLNO) resulting in graphs with higher likelihood scores and topological accuracy. The run with the highest log-likelihood per *m* value was selected.

### Test for introgression

Genotype probabilities from the SNP dataset used to infer migration events were used to calculate Patterson’s *D*, *f*_4_-ratio [34], and *f*-branch (*f*_b_) [33] statistics for all possible giraffe population trios using Dsuite v0.5-52 [35] with the okapi as an outgroup. The *f*_b_ is of particular interest as it can disentangle correlated *f*_4_-ratio estimates and assign evidence for introgression to specific, possibly internal, branches on a phylogeny given that they can be tested under a ((P1, P2) P3, Outgroup) topology. The admixture graph topology reconstructed by OrientAGraph, which was identical for *m* = 1 and *m* = 2, was used as the guide tree for the *f*_b_ estimation. Statistical significance was assessed through block-jackknifing with 100 equally sized blocks.

### Contemporary migration rates

Directionality and rates of contemporary migration between Nubian, reticulated, and Masai s. str. giraffe were estimated with BA3-SNPs v1.1 [79, 80]. First, a VCF file was generated by jointly calling genotypes in all individuals with median read depth ≥ 8 (Additional file 1: Table S1). Unlinked SNP sites identified in the second round of ANGSD were supplied to bcftools’ mpileup + call pipeline. Genotypes were called and sites were filtered as described for the SNP-based phylogenomic inference. We then randomly sampled ∼2% (8,137) of all SNPs using vcflib’s vcfrandomsample and converted the resulting VCF to the input format for BA3-SNPs with Stacks v2.59 [81] and stacksStr2immanc.pl (https://bit.ly/34fJdUz). Mixing parameters for BA3-SNPs were automatically tuned to achieve acceptance rates between 0.2 and 0.6 with BA3-SNPs-autotune v2.1.2 [82] by conducting short exploratory runs of 2.5 million iterations with a burn-in phase of 500,000 steps. The final BA3-SNPs run used 22 million iterations, a burn-in phase of 2 million steps, a sampling interval of 2,000 iterations, and the tuned mixing parameters (-m 0.1000 -a 0.2125 -f 0.0125). To assess chain convergence, the analysis was repeated three times, each starting from a different random seed, and the log probabilities of each run were plotted (Additional file 2: Fig. S8). The run with the smallest Bayesian deviance was selected [83] and the estimated migration rates were shown as a circular plot. We also constructed 95% credible sets using the formula *m* ± (1.96 × *sdev)*, where *m* is the posterior mean migration rate and *sdev* is the standard deviation of the marginal posterior distribution [79]. Migration rates were considered significant if the credible set did not include zero.

### Demographic reconstruction

Recent changes in *N_e_* of Nubian, reticulated, and Masai giraffe s. str. were assessed based on the SFS. First, a genome consensus sequence was generated for the okapi using ANGSD with option -doFasta 1 to polarize SNPs during the SFS estimation. We enabled -baq 2 and discarded sites with mapping or base qualities < 30 or depth below 4 or above the 95^th^ percentile of the sample’s depth distribution. We then calculated site allele frequencies per species in ANGSD with option -doSaf 1 and the okapi consensus sequence as ancestral. Individuals with median read depth < 2 were not included and quality filters were set as described for SNP calling. No HWE, MAF, and SNP *p*-value filters were used [84]. Site allele frequencies were converted into the unfolded SFS with ANGSD’s realSFS. Demographic histories were reconstructed from the unfolded SFS using Stairway Plot v2.1.1 [85], after masking singletons, with a mutation rate of 2.12 × 10^-8^ substitutions per site per generation estimated for the giraffe [86] and a generation length of 10 years [87].

## Supplementary Information

The online version contains supplementary material available at XXX.

**Additional file 1: Table S1.** Sample details and mapping statistics for analyzed individuals.

**Additional file 2: Figures S1–S8. Fig. S1.** Relatedness between pairs of individuals. **Fig. S2.** Principal component analysis (PCA). **Fig. S3.** Admixture analyses assuming a varying number of ancestry components (*K*). **Fig. S4.** Assessment of the fit of admixture models assuming K = 1–11 based on the correlation of residuals. **Fig. S5.** OptM output for the TreeMix runs with migration edges (*m*) ranging from 0 to 5. **Fig. S6.** Admixture graphs of giraffe populations and their corresponding residual fit. **Fig. S7.** Linkage disequilibrium (LD) decay in Nubian, reticulated, and Masai giraffe s. str. **Fig. S8.** Log probability trace and Bayesian deviance (D) for each BA3-SNPs run.

**Additional file 3: Table S2.** Posterior mean migration rates among Nubian, reticulated, and Masai giraffe s. str.

## Declarations

### Ethics approval and consent to participate

Sampling of giraffe skin biopsies was conducted under the appropriate access and research permits from the Kenyan authorities (#NEMA/AGR/109/2018/93, #KWS/BRM/5001, and #NACOSTI/P/18/50967/20704).

### Consent for publication

Not applicable.

### Availability of data and materials

Raw sequencing reads generated in this study are available at NCBI Short Read Archive under the BioProject accession PRJNA815626. Nucleotide sequences of mitochondrial genomes assembled in this study are available at GenBank under the accession numbers OM973995–OM974107. The code used to process and analyze the data generated in this study is available at GitHub (https://bit.ly/3LmwG4p).

### Authors’ contributions

Conceptualization, RTFC, SW, JF and AJ; methodology, RTFC; software, RTFC; validation, RTFC; formal analysis, RTFC; investigation, RTFC; resources, AM, SF, MO, DM, SM, JS-D, JF and AJ; data curation, RTFC; writing—original draft, RTFC, SW, JF and AJ; writing—review and editing, RTFC, SW, AM, SF, MO, DM, SM, JS-D, JF and AJ; visualization, RTFC; supervision, AJ; project administration, AJ; funding acquisition, SF, JF and AJ.

## Acknowledgements

We thank an array of partners, in particular government and NGO partners across Kenya who collaborated with and/or financially supported the Giraffe Conservation Foundation to permit, collect and include samples in this analysis, including Cleveland Metroparks Zoo, Governments of Botswana, Chad, Ethiopia, Kenya, Namibia, Niger, Tanzania, Uganda, and Zambia, Ivan Carter Wildlife Conservation Alliance, and San Diego Zoo Wildlife Alliance. We also thank Emma Vinson for her assistance in coding the R script used for relatedness filtering.

