## Additional file 2: Figures S1-S8 for "Genomic analysis reveals limited hybridization among three giraffe species in Kenya"

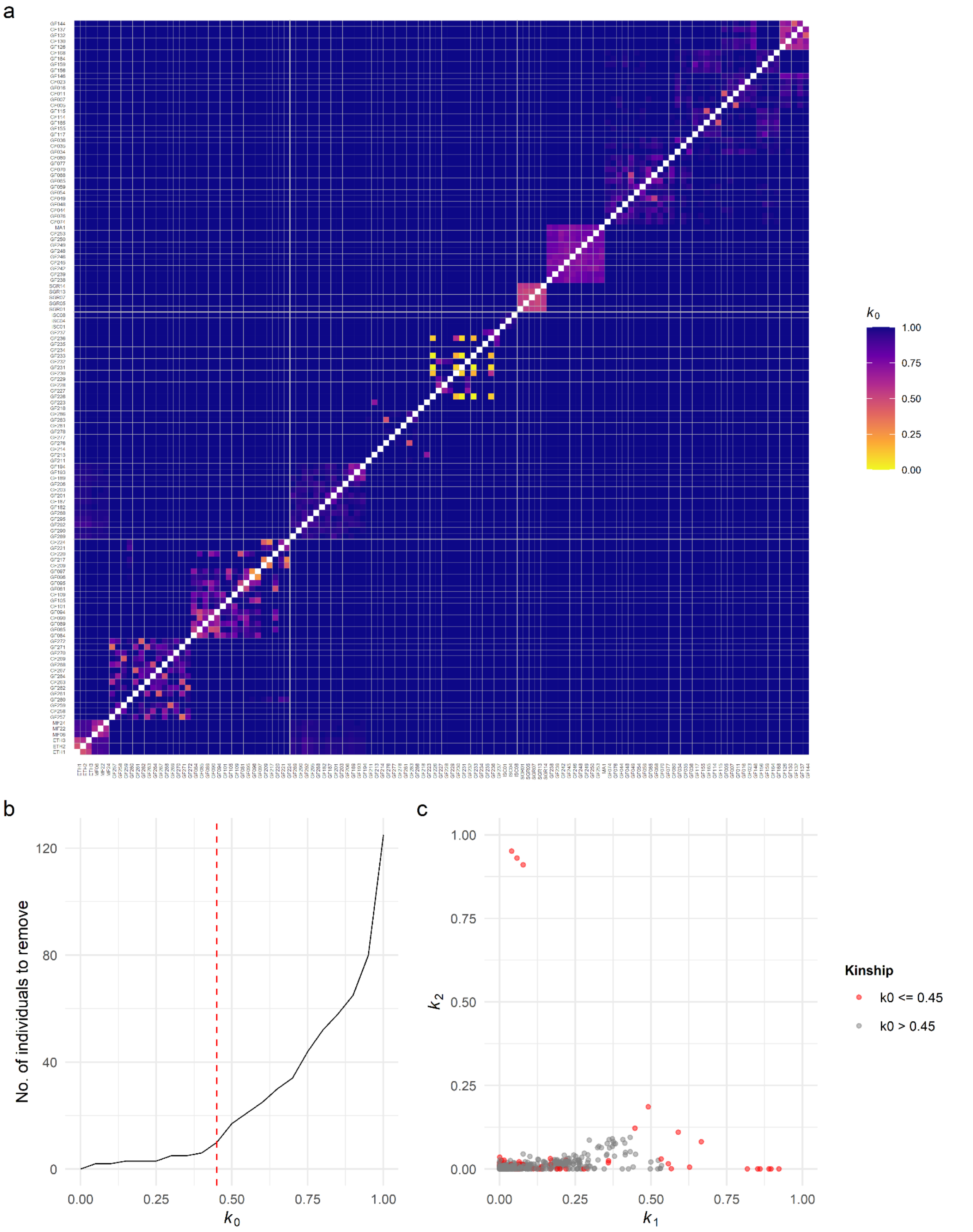


**Fig. S1. Relatedness between pairs of individuals.** (**a**) Pairwise relatedness coefficients estimated as $R$ = ($k_0$, $k_1$, $k_2$), where $k_0$, $k_1$, and $k_2$ are the proportion of the genome where a pair of individuals share 0, 1, or 2 alleles identical-by-descent, respectively. Warmer colors indicate higher relatedness. Individuals are sorted by taxon and sampling location as in Fig. 1c. (**b**) Number of individuals to remove assuming increasing thresholds for $k_0$. The red vertical line denotes the selected $k_0$ threshold (0.45) for relatedness filtering. Based on results from (a–b), individuals GF096, GF132, GF209, GF217, GF226, GF231, GF233, GF257, GF259, and GF262 were removed from further analyses. (**c**) Pairwise relatedness plotted as $k_1$ by $k_2$. Pairs of individuals affected by the filtering strategy ($k_0 \leq 0.45$) are shown in red.

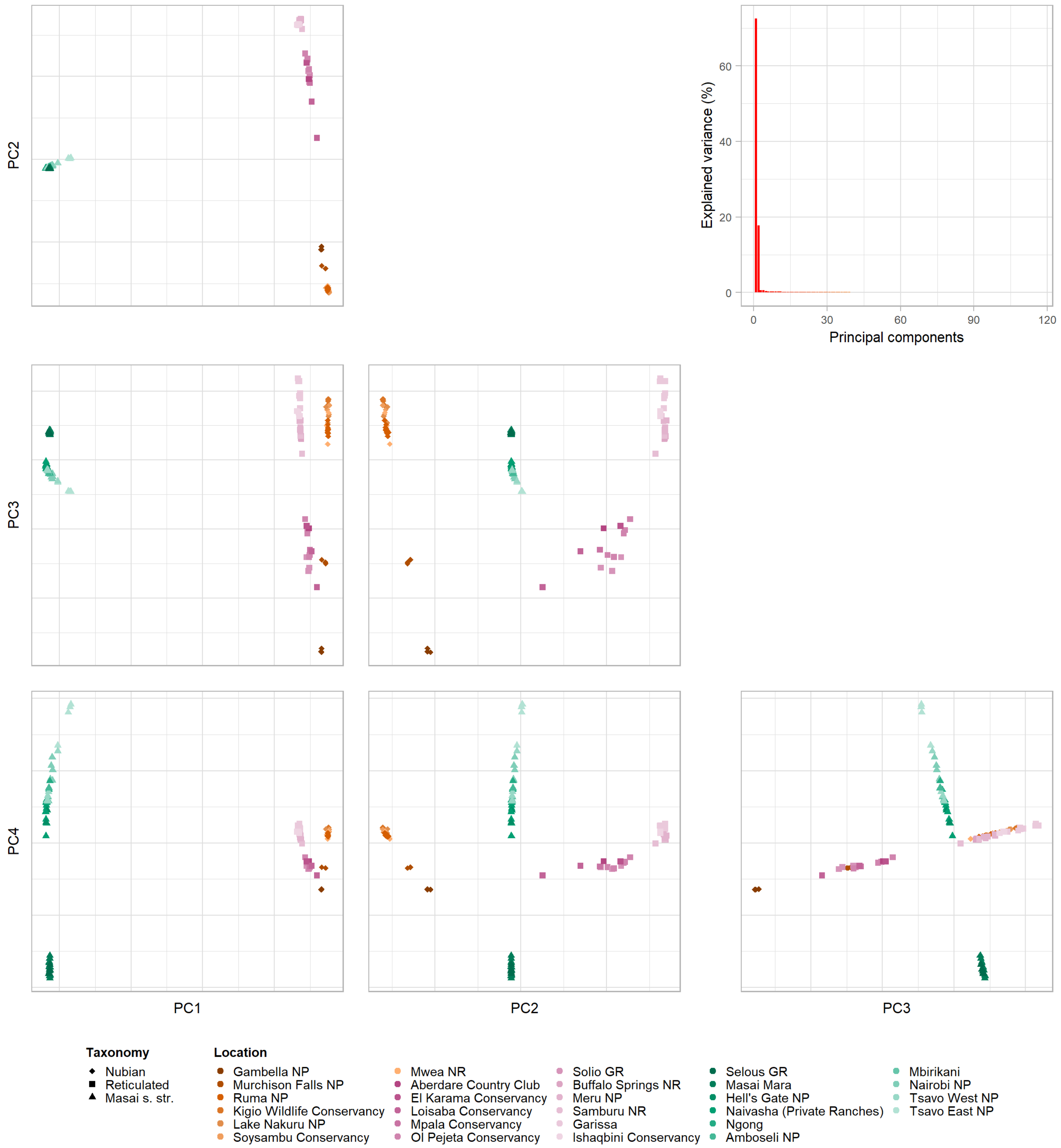


**Fig. S2. Principal component analysis (PCA).** PCA of 484,876 unlinked SNPs from 116 individuals representing Nubian, reticulated, and Masai giraffe s. str. Principal components (PCs) that explain at least 0.5% of the variance in the data are shown in a scatterplot matrix. Shapes represent the three focal taxa and colors represent sampling localities. In the top right is a scree plot showing the percentage of variance explained by each PC.

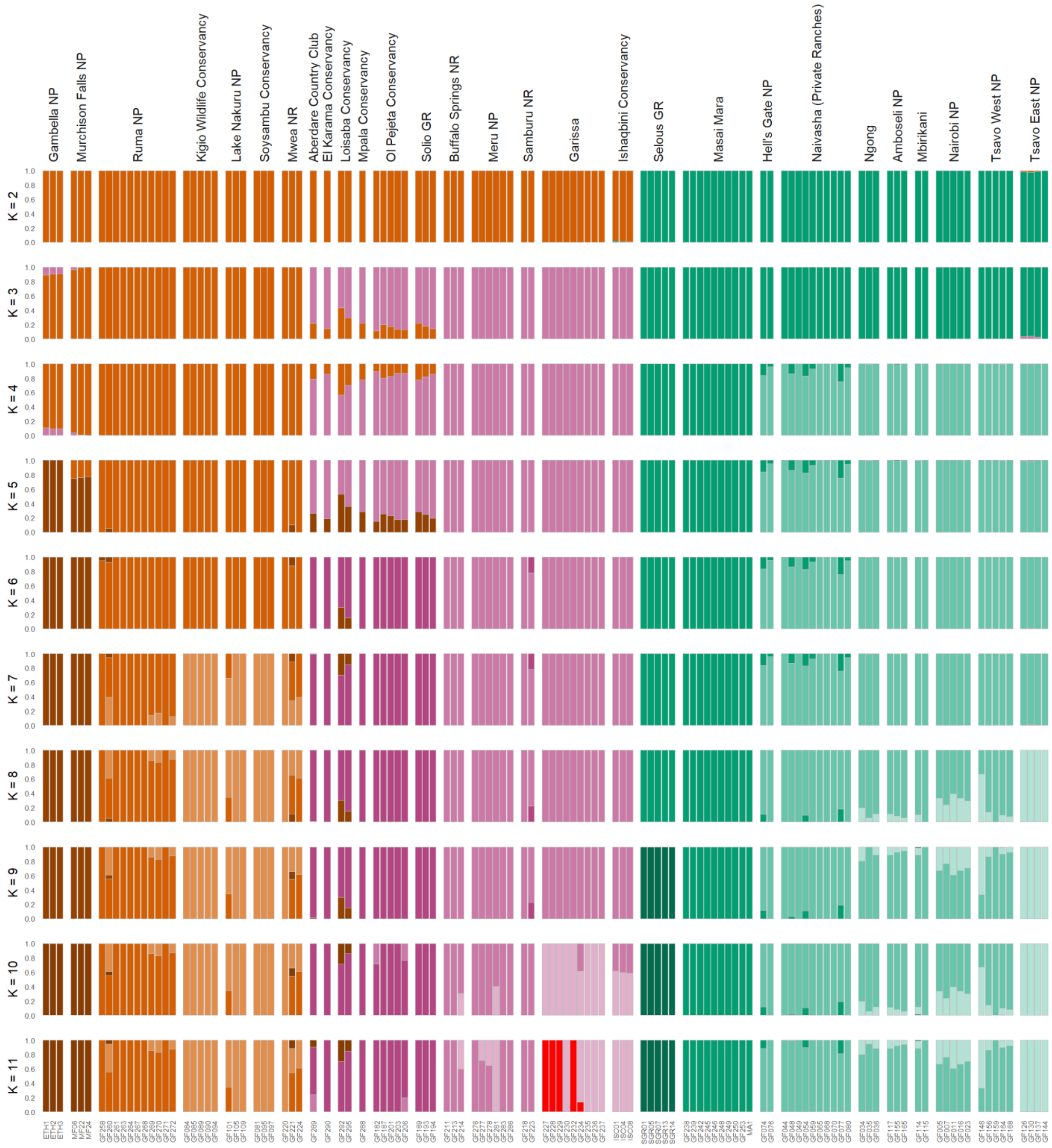


**Fig. S3. Admixture analyses assuming a varying number of ancestry components (*K*).** Ancestry proportions inferred from 484,876 unlinked SNPs from 116 individuals representing Nubian, reticulated, and Masai giraffe s. str assuming $K$ = 2–11. Individuals are sorted by sampling location. Colors indicate an individual's cluster membership.

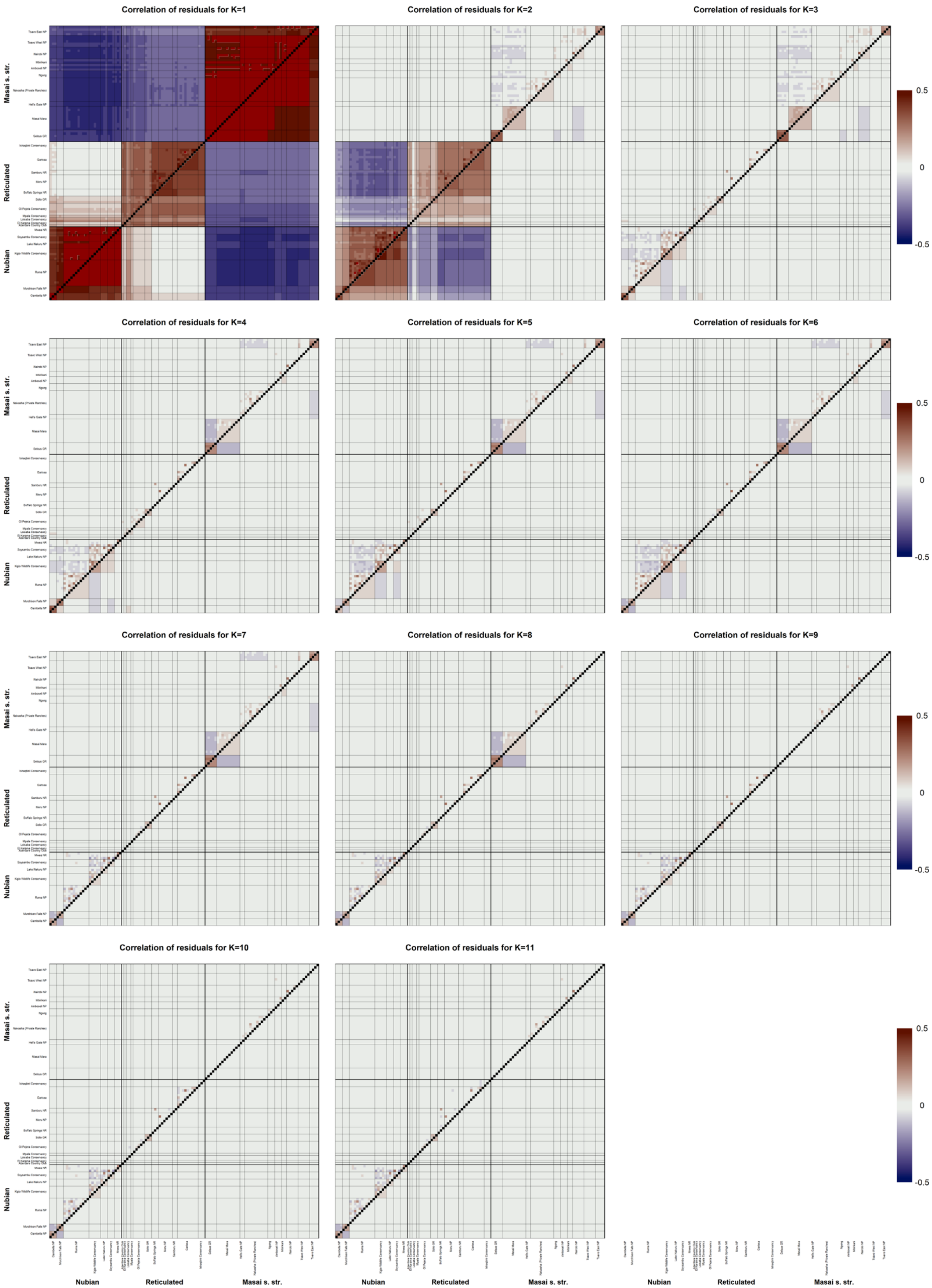

Correlation of residuals for K=1
Correlation of residuals for K=2
Correlation of residuals for K=3
Correlation of residuals for K=4
Correlation of residuals for K=5
Correlation of residuals for K=6
Correlation of residuals for K=7
Correlation of residuals for K=8
Correlation of residuals for K=9
Correlation of residuals for K=10
Correlation of residuals for K=11
Masai s. str.
Reticulated
Nubian
0.5
0
-0.5

**Fig. S4. Assessment of the fit of admixture models assuming *K* = 1–11 based on the correlation of residuals.** The assessed admixture models are shown in Fig. S3. The pairwise correlation of residuals between all individuals is shown above the diagonal, while the mean correlation within and between individuals from each sampling locality is plotted below the diagonal. The order of individuals and sampling localities are the same as in Fig. S3. Correlation values higher than 0.5 or lower than -0.5 are plotted as dark red and dark blue, respectively.

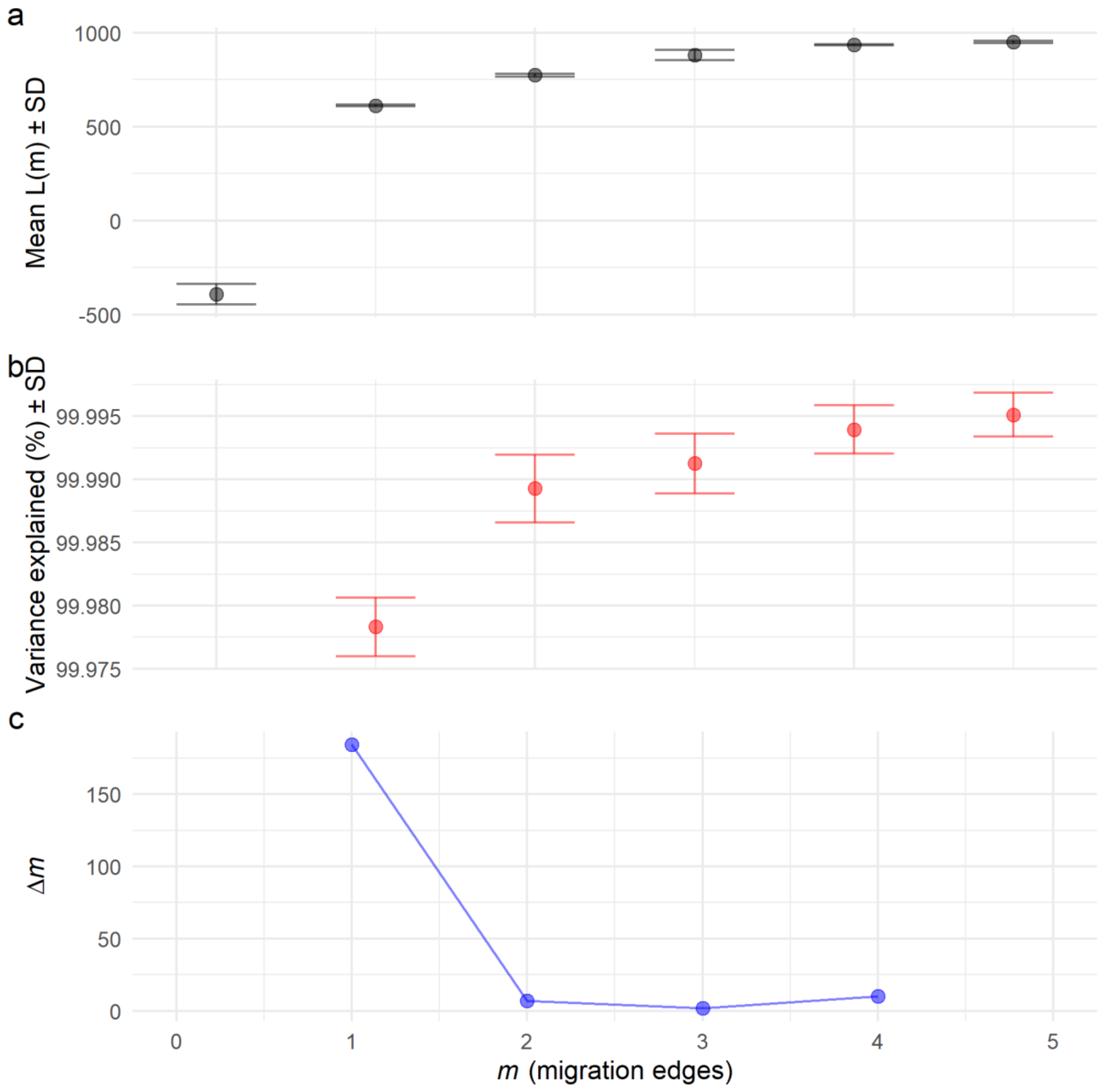


**Fig. S5. OptM output for the TreeMix runs with migration edges (*m*) ranging from 0 to 5.** (**a–b**) The mean and standard deviation (SD) across 50 iterations for the composite likelihood L(*m*) (black) and percentage of variance explained (red). (**c**) The second-order rate of change (*Δm*) across values of *m*.

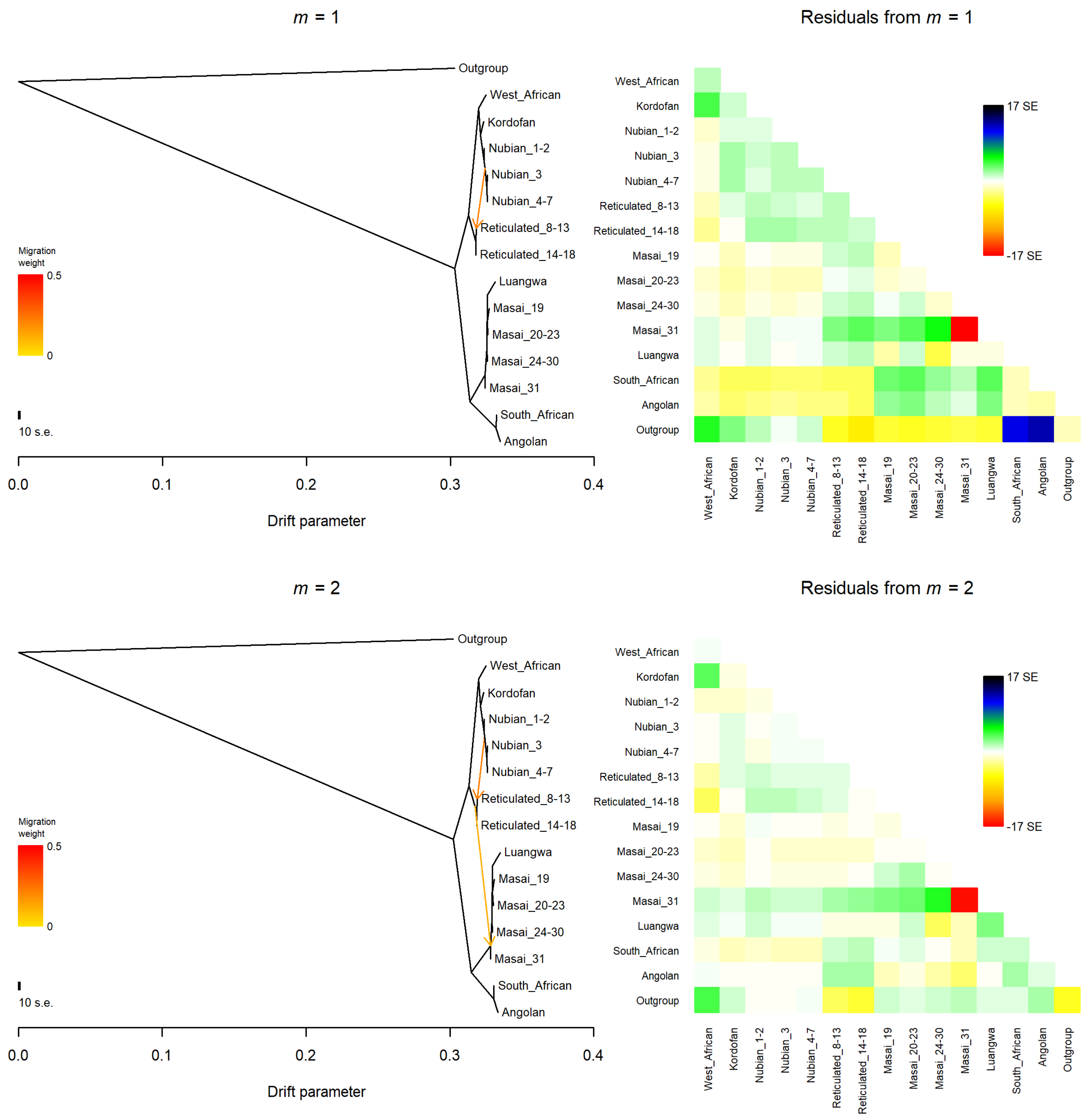


**Fig. S6. Admixture graphs of giraffe populations and their corresponding residual fit.** The graphs were estimated with OrientAGraph allowing for 1 and 2 migration events. The okapi was used as an outgroup. Nubian, reticulated, and Masai s. str. giraffe populations were defined following the best fit admixture model (*K* = 9; Fig. 1c–e, Fig. S3 and Fig. S4). Numerical suffixes in Nubian, reticulated, and Masai s. str. giraffe population labels indicate sampling locations according to Fig. 1a. Migration arrows are colored according to their weight. Positive residuals indicate populations that are more closely related in the dataset than in the best-fit graph, and thus are candidates for admixture events.

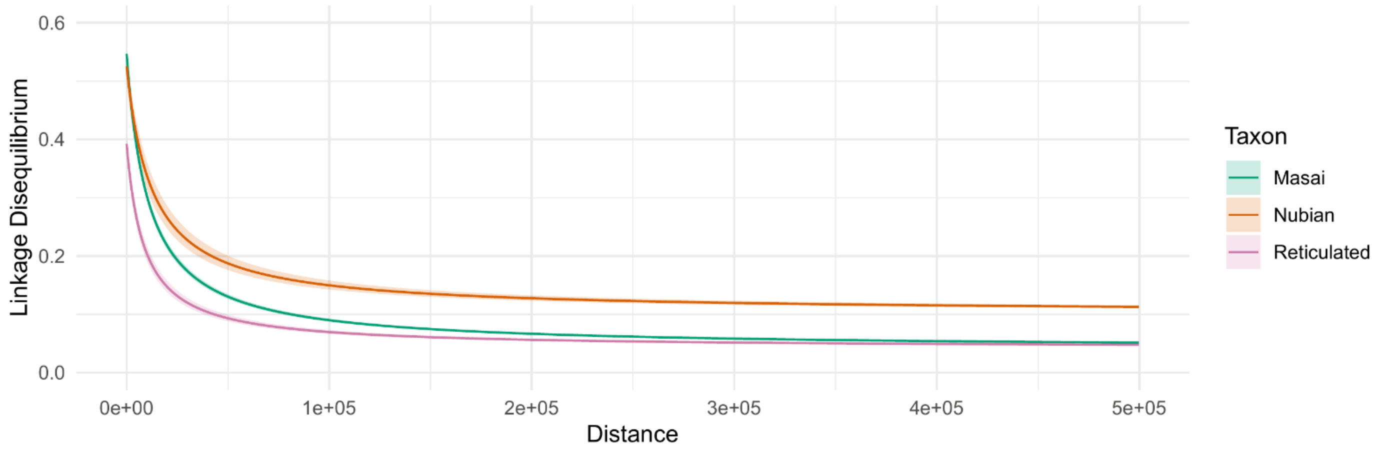


**Fig. S7. Linkage disequilibrium (LD) decay in Nubian, reticulated, and Masai giraffe s. str.** LD was estimated as $r^2$ values for SNP pairs up to 500 kbp apart. Linkage decay curves were fitted per species from a random sample of 0.05% of the estimated $r^2$ values.

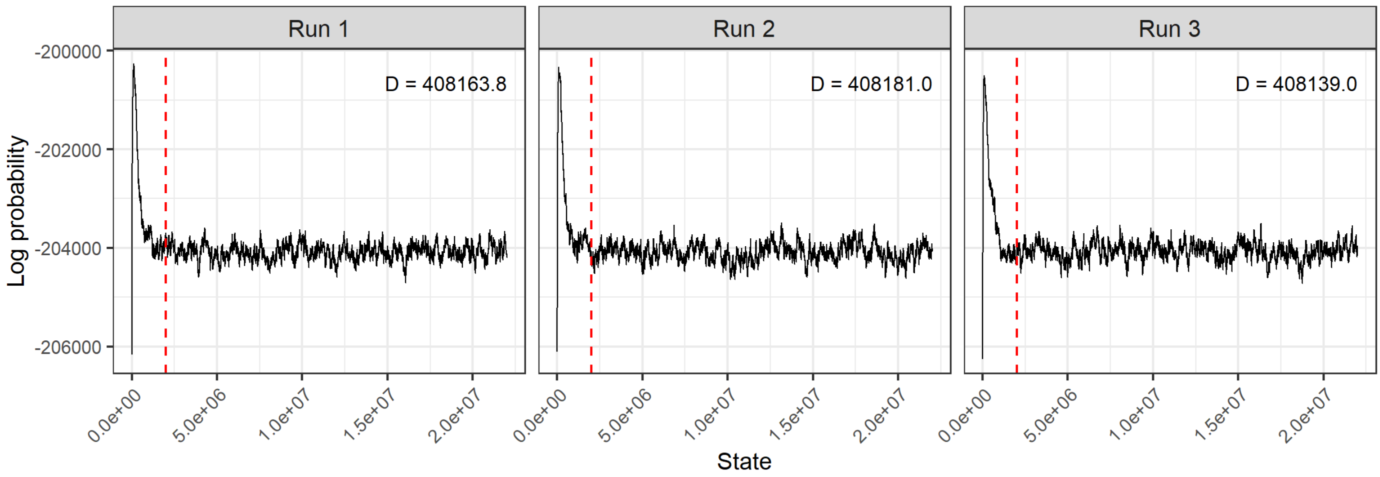


**Fig. S8. Log probability trace and Bayesian deviance (D) for each BA3-SNPs run.** The stationary log probability trace after the burn-in period (dashed vertical line) suggests convergence between runs. Run 3 shows the smallest D and was selected for reporting estimated migration rates on Fig. 3c and Additional file 3: Table S2.
